# Disruption of glutamine carrier *Slc38a1* causes cognitive impairment, anxiety and depressive-like behavior

**DOI:** 10.64898/2026.05.20.726495

**Authors:** Zubeide Sleeman, Gezime Seferi, Prabhat Khanal, Knut Tomas Dalen, Cecilie Morland, Farrukh Abbas Chaudhry

## Abstract

GABAergic deficit is associated with key neuropsychiatric disorders, such as major depressive disorder (MDD), anxiety, schizophrenia, and autism spectrum disorder (ASD). However, it is not known whether these disorders are causal to or a result of GABAergic dysfunction. We previously showed that the Solute carrier 38 member 1 (Slc38a1) accumulates glutamine in subpopulations of GABAergic neurons and sustains neurotransmitter GABA synthesis. Genetic inactivation of *Slc38a1* in mice caused lowered GABA levels, altered synaptic vesicle morphology, slowed γ-oscillations, and reduced cortical processing and plasticity, selectively at GABAergic synapses. We now demonstrate a significant reduction in learning and memory performance in the Morris water maze and increased signs of despair in the forced swim test in *Slc38a1*^-/-^ mice compared to *Slc38a1*^+/+^ mice, implicating cognitive impairments and depressive-like behavior. Examination in the open field maze also indicates anxiety and/or reduced interest in exploration. There are no signs of impaired sociability or recognition of social novelty in the three-chambered test, speaking against involvement in schizophrenia- or ASD-like disorders. Metabolic phenotyping and measurement of the locomotion do not segregate the *Slc38a1* genotypes, suggesting that the cognitive impairments, depressive-like behavior and anxiety are brain-dependent. Our data is further supported by a pathologic variant of *Slc38a1* in a family with depression and suicidal behavior. Altogether, we demonstrate that dysfunction of Slc38a1-dependent GABA synthesis and the ensuing impaired γ-oscillations underpin the pathogenesis of neurocognitive deficits, anxiety and depression.

## INTRODUCTION

Neuropsychiatric and neurological disorders represent a substantial and growing global health burden. Despite their diverse clinical manifestations, these conditions converge on a common feature: disrupted inhibitory neurotransmission. Central to this, GABA (γ-aminobutyric acid) signaling regulates neural circuit stability and cognitive function, yet the mechanisms linking its dysregulation to disease pathogenesis remain incompletely understood. GABA is the principal inhibitory neurotransmitter and contributes to the fine-tuning of synaptic plasticity, action potential firing, synchronization of neural networks, neurodevelopment, and the excitatory/inhibitory balance in the brain^1,2^. GABA signaling is fundamental for normal cognitive function, and abnormality underpins main neuropsychiatric diseases, such as MDD, anxiety, schizophrenia, ASD, Alzheimer’s disease, and epilepsy^3,4,5^. Several of these conditions co-cluster and have common genetic risk factors^6,7^, and their pathophysiology is associated with reduced activities of the GABA-synthesizing enzyme glutamic acid decarboxylase 67 (GAD67), GABA receptors, parvalbumin-positive (PV^+^)-, and/or somatostatin-positive (SST^+^)-interneurons^8,9,10^. However, whether these changes drive neuropsychiatric diseases or are consequences of them, has not been resolved.

PV^+^-interneurons and the associated GABAergic system are highly vulnerable to stressors and may be core factors leading to neuropsychiatric diseases^5,11^. PV^+^-interneurons are interconnected through chemical and electrical synapses and synchronize neuronal firing^12^. In addition, they are intrinsic for generating cortical γ-oscillations, which increase cortical circuit performance involved in higher cognitive functions, such as working memory, storage and retrieval of information from long-term memory, and attention^13,14,15,11^. Dysfunction of γ-oscillations associated with PV^+^-interneuron is a hallmark of several neuropsychiatric disorders^16^. Despite the extensive literature supporting the involvement of dysfunctional GABA systems and the PV^+^-interneurons in neuropsychiatric diseases, the underlying GABAergic deficits and/or pathological variants of the proteins involved are yet to be revealed mechanistically^17^. We have previously molecularly identified and characterized the System A transporter Slc38a1 and we have shown that it translocates neutral amino acids across cell membrane^18^. Slc38a1 is enriched in PV^+^- and SST^+^-interneurons and preferentially transports glutamine unidirectionally into these cells^19,20,21^. The Slc38a1-mediated intracellular accumulation of glutamine is pivotal for the synthesis of the neurotransmitter GABA^22^. In addition, translocation of glutamine through the Slc38a1 transporter also induces high-frequency membrane oscillations, which are able to generate long-lasting action potential^23^. Genetic inactivation of *Slc38a1* in mice waned GABA synthesis and levels of vesicular GABA, and altered vesicular morphology despite a significant compensatory upregulation of GAD67 and phosphate-activated glutaminase (PAG) to increase the synthesis of GABA^23^. Moreover, disruption of *Slc38a1* modifies PV^+^-interneuron dependent γ-oscillations and impairs cortical processing and plasticity. In the current study, we have determined the behavioral phenotype of Slc38a1 null-mutant (*Slc38a1*^-/-^), heterozygous mutant (*Slc38a1*^+/-^) and wild type (wt; *Slc38a1*^+/+^) mice by measuring the impact of *Slc38a1* inactivation on a number of neuropsychiatric functions, metabolism, and locomotion. Our data demonstrates that Slc38a1 dysfunction may induce cognitive impairment, depressive-like behavior and anxiety, but not the pathogenesis of schizophrenia and ASD.

## MATERIAL AND METHODS

A summary of the main material and methods are described below. Detailed information on material and methods is elaborated in Supplementary information.

### Animals

This study was conducted in strict accordance with the recommendations described in the Norwegian Animal Welfare Act and the European Union Directive on the Protection of Animals used for Scientific Purposes (2010/63/EU). All experiments have been approved by the Norwegian Food Safety Authority (FOTS application numbers 9200, 10902, 21009 and 30120) and reported in accordance with reporting of In Vivo Experiments (ARRIVE)^24^. Mice were housed in a room with fixed 12-hour light (7 am-7 pm)/dark (7 pm-7 am) cycle, temperature set to 22 ± 1 °C with relative humidity at 50 ± 10%. The animals were stalled in groups up to 9 mice in Makrolon GreenLine cages (Sealsafe Plus GM500 or GM900, Buguggiate, Italy). The cages were always ventilated with 100% fresh air except during experiments. The cages were enriched with paper for nest-building, a wooden stick, a paper roll or a small plastic house and a running wheel. Autoclaved food (RM3 from Special Diets Service (UK)) and water were provided *ad libitum*.

The molecular impact of *Slc38a1*^-/-^ mouse model used in these studies has previously been rigorously characterized^23,22^. Animals used for the study were backcrossed into C57BL/6J for 10 generations. Mice that were wild type (*Slc38a1*^+/+^), heterozygous for the mutation (*Slc38a1*^+/-^), or genetically inactivated for *Slc38a1* (*Slc38a1*^-/-^) were all originating from the same breeding colony. Animals included in the study were of both sexes and aged 10-15 weeks, except for metabolic phenotyping where they were 22-23 weeks. A total number of 117 animals were used in this project, divided in five groups consisting of respectively 23, 26, 14, 27 and 27 animals. Cages of several litters were used. Before each behavioral test all animals had a habituation period of one hour to acclimate to the experimental environment. All behavioral tests were performed in the same room, at the same time of the day (between 1:00 p.m. and 6:00 p.m.) and by the same experimenter to minimize any effects of environmental factors. The smell of male mice can make female mice anxious, therefore, experiments were first performed on all female mice followed by the male mice.

### Morris water maze

The Morris water maze setup consisted of a circular pool that was placed in a room with white surroundings and high-contrast visual cues on each wall. The pool was filled with opaque water and the arena was digitally divided into four quadrants (north, south, east and west) by an advanced video tracking system (ANY-maze; Stoelting Europe, Ireland).

The tests were performed using a 6-day protocol. A white platform was placed in the northern quadrant and submerged 0,5 cm below the water surface throughout the learning trials (day 1-5). Test animals were trained to learn the position of the platform by identifying distal visual cues. For each learning day, all animals were released at four different locations. The animals were then given 60 seconds to swim and search for the platform, followed by 30 seconds on the platform.

A probe test was performed on day 6 with a hidden platform. All animals were released in a new start position. The probe test consisted of only one trial and animals were allowed to swim freely for 60 seconds. 11 animals were excluded because they floated for more than 30 seconds, this criteria was set a priori.

### Forced swim test

During the forced swim test each animal was placed in its own glass cylinder filled with clear water. The experimenter observed the mice via a GoPro^TM^ camera to ensure that the test animals never appeared to be in serious distress, for a six-minute time lapse. However, only the last four minutes were analyzed due to the disturbances on measurements caused by excessive activity at the beginning^25^. The experimenter used a stopwatch to measure mobility time to state the immobility time. The definition of mobility time is “*any movements other than those necessary to balance the body and keep the head above the water”*^25^. The experimenter was trained to correctly identify movements that are counted as mobility. In addition, the experimenter was blind to the genotype and recordings were analyzed twice to minimize bias.

### Elevated plus maze

The elevated plus maze apparatus was shaped as a plus sign with two facing closed arms and two facing open arms. The animal was placed at the center zone and was then allowed to explore the maze freely for five minutes. Each mouse got only one test trial. A video tracking camera placed above the apparatus was connected to ANY-maze that automatically collected data.

### Open field test

The open field test was performed in a VersaMax Animal activity monitor (AccuScan Instruments, Inc., USA). The mice were placed in the center of the maze and were allowed to explore it freely for 60 minutes. Mice movements were tracked for all 60 minutes by infrared sensors in the floor of the square activity monitor. The arena was digitally divided into a center zone and a peripheral zone. One animal was excluded when testing for normal distribution.

### The three-chambered social test

The three-chambered sociability test setup was divided by removable partitions with openings to allow the animals to move freely. Each of the distal chambers contained a wired cage. This experiment consists of a habituation period, the sociability test and a preference for social novelty test. During the habituation period the test mouse was allowed to explore the apparatus for 10 minutes with empty wired cages. For the sociability test, a C57Bl/6J mouse (stranger 1) which was unfamiliar to the test mouse was placed in one of the wired cages. For the preference for social novelty test, a new stranger mouse (stranger 2) was placed in the previously empty wired cage, and the test mouse was allowed to move freely for another 10 minutes. The experiment was videotaped and connected to ANY-maze.

### The notched beam test

The notch beam test apparatus was a 1-meter-long beam consisting of blocks with gaps in between. Mice were trained to cross the beam three times a day for three consecutive days, without stalling. On the test day, the mice were videorecorded with a GoPro^TM^ camera. Time spent to transverse the center 50 cm of the beam was measured by a stopwatch, and the number of paw slips (defined as the foot coming off the top of the beam^26^) was counted by analyzing the video recordings. Two successful trials in which the mouse did not stall on the beam were averaged and further analyzed. 3 animals were excluded because they obviously stalled on test day.

### Metabolic phenotyping

Male and female mice were placed in a metabolic cage system consisting of several cages (Phenomaster, TSE Systems, Germany). After 36-40 hours of single housing and acclimatization in the metabolic cages, indirect calorimetric measurement of oxygen consumption and carbon dioxide production were performed. Physical activity was measured as movement in the XY-plane^27^. Data collected over a 48-hour period were used to calculate mean values in the light (12 hours) and the dark (12 hours) phases, respectively. Respiratory exchange ratio (RER) values were calculated based on measured O_2_ and CO_2_ values. Body weights were measured prior to metabolic phenotyping.

### Statistics

All results were tested for normal distribution visually with qq-plots before analyzing further. Either One-way ANOVA or Two-way ANOVA followed by post hoc Tukey’s test was performed to compare different parameters between *Slc38a1*^+/+^, *Slc38a1*^+/-^ and *Slc38a1*^-/-^ mice, except for the notched beam test where unpaired t-test was used to compare *Slc38a1*^+/+^ and *Slc38a1*^-/-^ mice. All statistical analyses were performed by GraphPad Prism Version 9.0.

## RESULTS

### *Slc38a1*^-/-^ mice display spatial learning and memory impairment

Behavioral phenotyping may disclose the impact of a gene variant on cognitive skills, psychiatric disorders, metabolism and more^28^. We, therefore, investigated cognitive functions in the three *Slc38a1* mouse variants by Morris water maze, a behavioral test for hippocampal-dependent learning and memory. The test measures the ability of mice to learn and remember the location of a hidden platform in a circular pool based on distal clues^29^.

In the Morris water maze, the latency to escape to the platform was quantified for 5 consecutive training days followed by a probe test on the 6^th^ day. *Slc38a1*^+/+^ mice show a significant improvement from day 1 to day 4 and 5 (p<0.05 and p<0.0001, respectively) and from day 2 and 3 to day 5 (p<0.0001 and p<0.01 respectively; Fig. 1A). The *Slc38a1*^+/-^ mice reach significant acquisition only between day 1 and day 5 (p<0.01). In contrast, *Slc38a1*^-/-^ mice do not improve escape latency to the platform from day 1 to day 5 and show significantly longer escape latency compared to *Slc38a1*^+/+^ mice on day 5 (Fig. 1A). Altogether, the steep escape latency curve for *Slc38a1*^+/+^ mice suggests fast task acquisition, while *Slc38a1*^-/-^ mice have flat learning curve with no significant reduction between day 1 and day 5 implicating a deficit in task acquisition (Fig. 1A). *Slc38a1*^+/-^ mice have a shallower curve consistent with slower task acquisition compared to *Slc38a1*^+/+^ mice.

**Figure 1.**
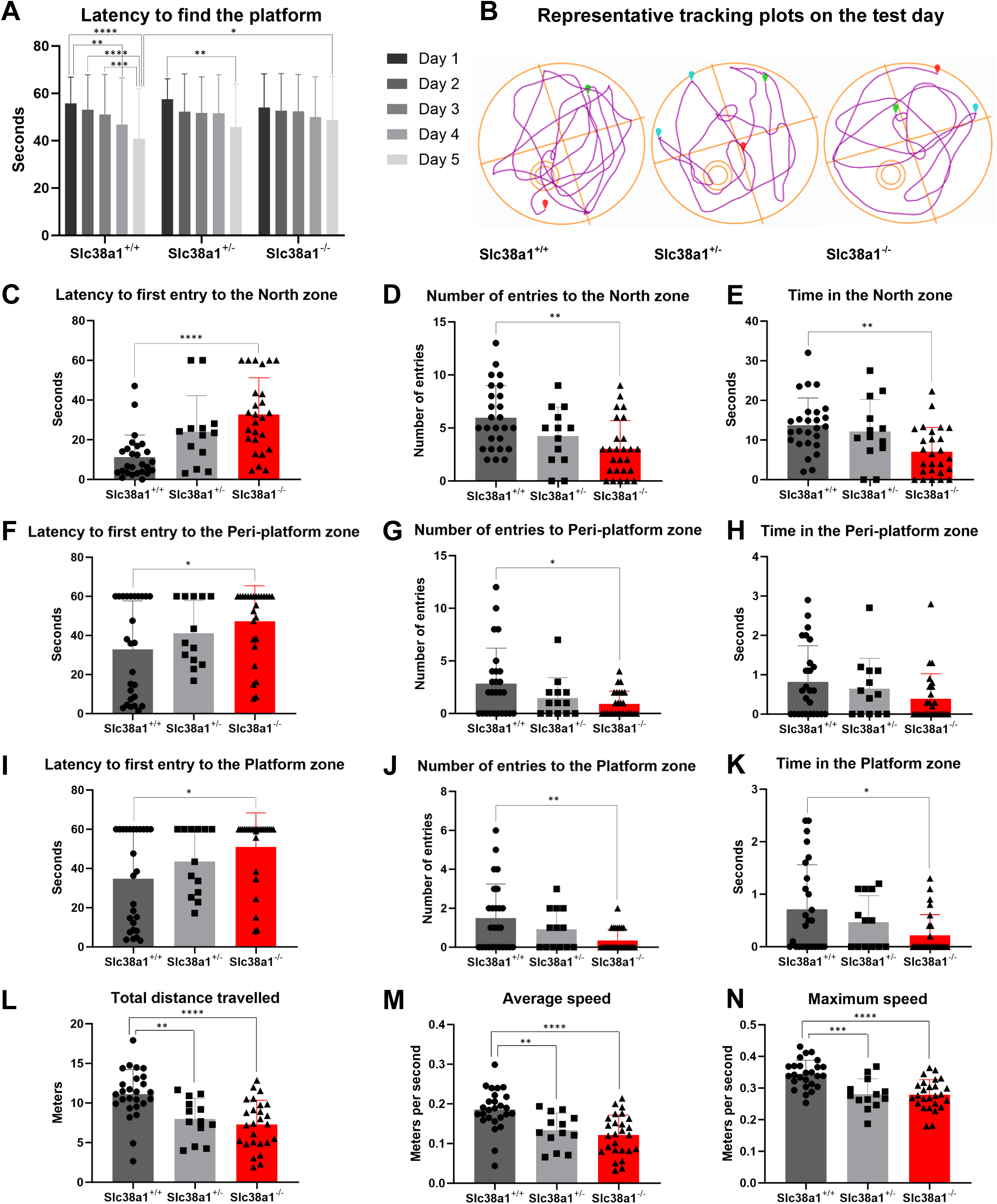
Genetic inactivation of *Slc38a1* in mice reduces spatial learning and reference memory in Morris water maze. Spatial learning and memory in *Slc38a1*^+/+^, *Slc38a1*^+/-^ and *Slc38a1*^-/-^ mice were tested in a Morris water maze consisting of a white circular pool filled with opaque water with or without a platform placed under the water surface. (**A**) During the first learning day of including platform tests, the *Slc38a1*^+/+^, *Slc38a1*^+/-^ and *Slc38a1*^-/-^ mice exhibit a similar latency to escape onto the visible platform. In learning trial days including the platform, *Slc38a1*^+/+^ mice show a significantly shorter latency to escape onto the platform on day 4 compared to day 1 and on day 5 compared to day 1, 2 and 3. The *Slc38a1*^+/-^ mice show a shorter latency to escape on day 5 compared to day 1. The *Slc38a1*^-/-^ mice showed no significant difference in latency on the learning days including the platform and latency to escape on day 5 is significantly longer compared to *Slc38a1*^+/+^ on day 5. (**B**) The tracking plots show representative swimming patterns of *Slc38a1*^+/+^, *Slc38a1*^+/-^ and *Slc38a1*^+/+^ mice, respectively, in the Morris water maze on probe trial day without a platform. Green marks “test start”, red marks “test end” and blue marks “freezing”. (**C-E**) On probe trial without a platform on day 6, the *Slc38a1*^+/+^ mice show a significantly shorter latency to the first entry to the north zone (where the platform was located during learning days), a higher number of entries and more time spent in the north zone compared to the *Slc38a1*^-/-^ mice. The swimming pattern of *Slc38a1*^+/-^ mice reflect a hybrid pattern of *Slc38a1*^+/+^ and *Slc38a1*^-/-^ mice and do not show any significant changes compared to the other two genotypes. (**F-H**) The peri-platform zone is 3 cm wider than the platform zone. The *Slc38a1*^+/+^ mice find the peri-platform zone significantly faster than the *Slc38a1*^-/-^ mice in addition to entering this zone more frequently on the probe trial. The *Slc38a1*^+/-^ mice perform in between the two other genotypes and do not deviate significantly from them. (**I-K**) On probe trial, the latency of first entry to the platform zone is lower, and the number of entries and time spent there is significantly higher in *Slc38a1*^+/+^ mice compared to *Slc38a1*^-/-^ mice. *Slc38a1*^+/-^ mice perform in between the *Slc38a1*^+/+^ and *Slc38a1*^-/-^ mice but do not show significant differences to either of the genotypes. (**L-N**) During the probe trial, *Slc38a1*^+/+^ mice travel significantly larger total distance and with significantly higher average and maximum speed compared to *Slc38a1*^+/-^ and *Slc38a1*^-/-^ mice. In A and C-N, bars represent mean±SD for *Slc38a1*^+/+^ (dark grey), *Slc38a1*^+/-^(grey) and *Slc38a1*^+/+^ (red). Asterisks indicate the level of significance by two-way ANOVA (B) / one-way ANOVA with post hoc Tukey’s test (*p < 0.05, **p < 0.01, ***p < 0.001, ****p < 0.0001). 26 *Slc38a1*^+/+^, 13 *Slc38a1*^+/-^ and 26 *Slc38a1*^-/-^ mice were used in this experiment.

On the test day (day 6), the tracking plots indicate that *Slc38a1*^+/-^ and *Slc38a1*^-/-^ mice targeted the platform area less frequently and had lower mobility than *Slc38a1*^+/+^ mice (Fig. 1B). We next calculated latency to first entry, number of entries during the 60 seconds of the trial, and time spent in three zones in the pool, i.e., the north zone (the quadrant where the platform was located), the peri-platform zone (3 cm wider than the platform) and the platform zone itself. *Slc38a1*^-/-^ mice display a substantially longer latency to the first entry to the North zone compared to *Slc38a1*^+/+^ mice (p<0.0001; Fig. 1C). Accompanying this, the *Slc38a1*^-/-^ mice had significantly lower number of entries and time spent in the North zone (p<0.01; Fig. 1D, E). Upon narrowing the area of interest to the peri-platform zone or the platform zone, the same pattern as for the entire North zone was observed: *Slc38a1*^-/-^ mice show an increase in the latency to first entry and a reduction in the number of entries to both zones comparted to their *Slc38a1*^+/+^ littermates (Fig. 1F-G, I-J). Furthermore, there was a reduction in the time spent in both zones, but this only reached statistical significance for the platform zone (Fig. 1H, K). For all the parameters mentioned above, the *Slc38a1*^+/-^ mice displayed intermediate values between *Slc38a1*^+/+^ and the *Slc38a1*^-/-^ mice, but these values were statistically indistinguishable from both of the other genotypes (Fig. 1F-N). *Slc38a1*^-/-^and *Slc38a1*^+/-^ mice were less mobile than *Slc38a1*^+/+^ with significantly lower total distance traveled as well as lower average and maximum speed (Fig. 1L-N). Finally, the results were investigated for interaction by sex, but there were no significant differences in results between males and females (Fig. S1).

### *Slc38a1*^-/-^ mice display depressive-like behavior

We next performed a forced swim test to investigate for depressive-like behavior of these mice^30^. *Slc38a1*^-/-^ mice exhibit a small, but statistically significant increase in immobility time compared to *Slc38a1*^+/+^ mice (Fig. 2A). Again, the *Slc38a1*^+/-^ mice showed intermediate immobility times, statistically indistinguishable from both other genotypes. Stratifying the results into males and females showed no differences between the two sexes (Fig. 2B). Overall, the increase in immobility time in *Slc38a1*^-/-^mice suggests that impairment of GABA signaling causes depressive-like behavior.

**Figure 2.**
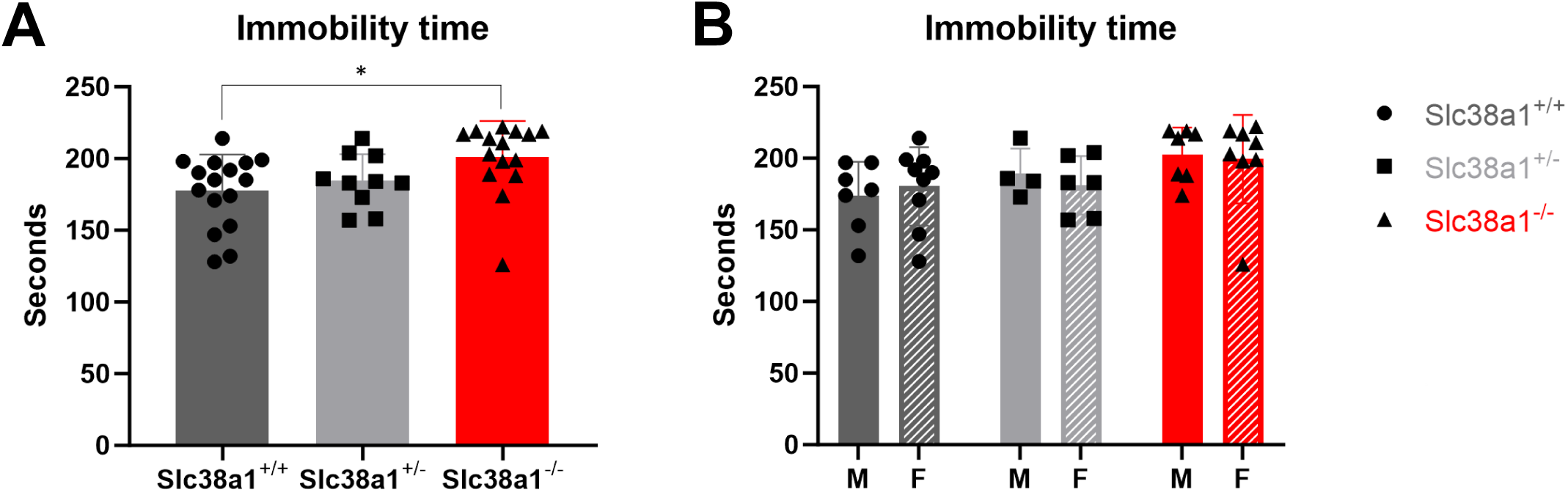
*Slc38a1*^-/-^ are more immobile compared to *Slc38a1*^+/+^ in the forced swim test. The mobility of test animals was investigated by the forced swim test as a proxy for depressive behavior, where mice were placed in containers filled with water and their behavior was observed. (**A**) *Slc38a1*^-/-^ mice exhibit a significantly more immobile behavior than *Slc38a1*^+/+^ mice. (**B**) No significant difference in immobility time was seen between males and females within the three genotypes. The bars show mean±SD, with dark grey representing *Slc38a1*^+/+^, grey *Slc38a1*^+/-^ and red *Slc38a1*^-/-^ mice (respectively). In (**B**) solid bars represent males and hatched bars represent females. The data are analyzed by one-way ANOVA (A) / two-way ANOVA (B) with post hoc Tukey’s test (*p < 0.05). 7 males and 9 females of *Slc38a1*^+/+^, 4 males and 6 females of *Slc38a1*^+/-^ and 7 males and 8 females for *Slc38a1*^-/-^ mice were included.

### *Slc38a1*^-/-^ mice display signs of anxiety and/or lack of interest in exploration

Anxiety-like behavior in the three *Slc38a1* genotypes was measured in the elevated plus maze, which allows testing the natural aversion of mice for open and elevated areas and their natural spontaneous exploratory behavior in novel environments^31^. Track plots show that mice of all three *Slc38a1* variants explore both closed and open arms of the elevated plus maze, however, lowered overall mobility of the *Slc38a1*^-/-^ mice was observed compared to *Slc38a1*^+/+^ littermates (Fig. 3A). Indeed, the total distance travelled within the test time was significantly less for *Slc38a1*^-/-^ mice compared to *Slc38a1*^+/+^ mice (p<0.01; Fig. 3B). The *Slc38a1*^-/-^ mice enter both arms and the center less frequent, and spend less time in the center, compared to *Slc38a1*^+/+^ and *Slc38a1*^+/-^ mice (Fig. S2). We then tested whether the mice prefer the open arms, closed arms, or the center area. Our data show that the three *Slc38a1* genotypes have equal preference for the entries into open arms and the closed arms (Fig. 3C-E). *Slc38a1^+/+^* mice also spend significantly more time in the closed arms than in the open arms, as is expected for this strain, and this does not differ in *Slc38a1*^-/-^ mice (Fig. 3F-H).

**Figure 3.**
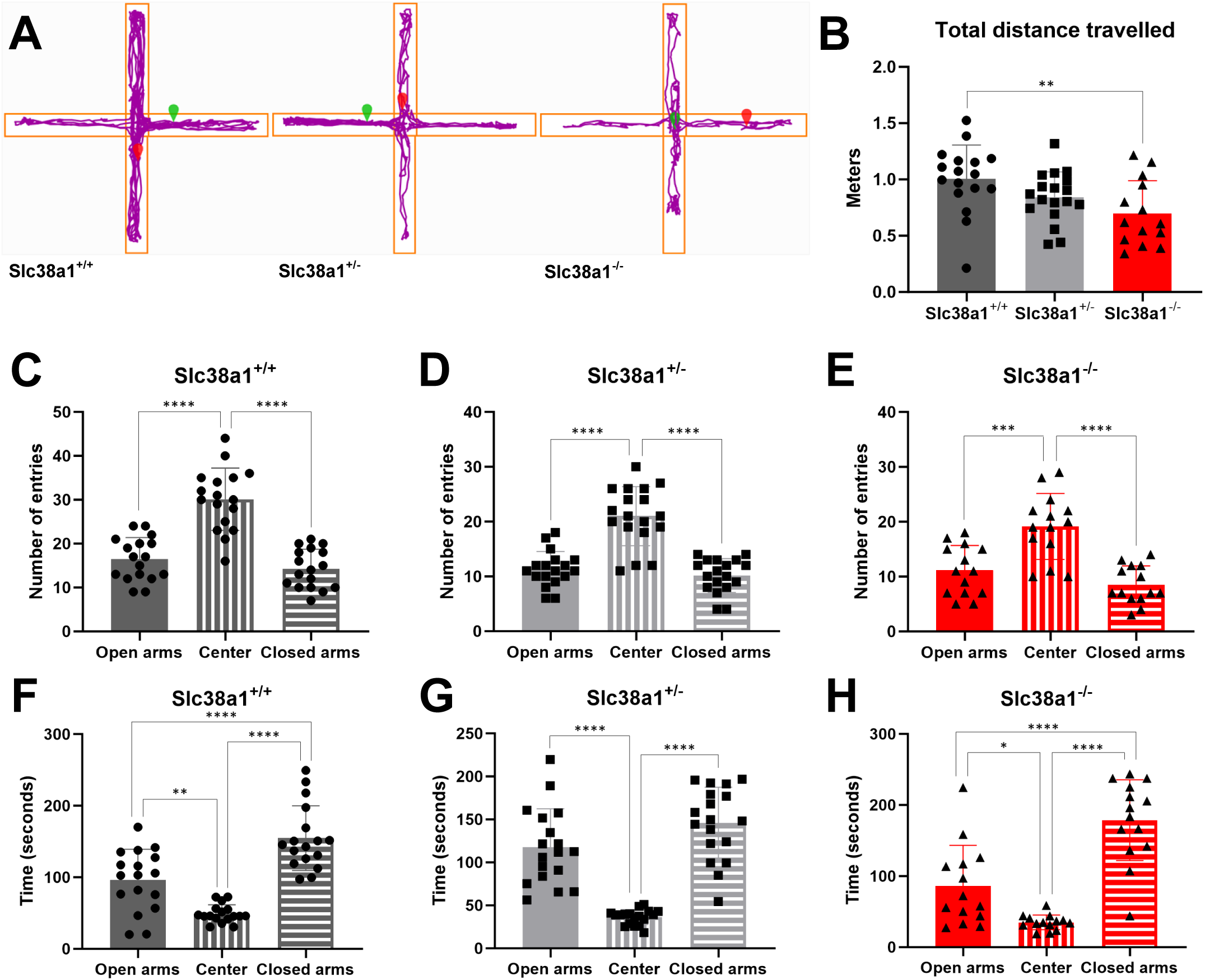
Wild type mice and mice impaired for *Slc38a1* show no mutual differences in preference for open or closed arms in the elevated plus maze. Natural spontaneous exploratory and anxiety-like behavior were investigated in an elevated plus maze configurated as a plus (+) with two closed arms (horizontal) perpendicular to the open arms (vertical), and a center platform. (**A**) Representative tracking plots show that *Slc38a1*^-/-^ and *Slc38a1*^+/-^ mice are less active than *Slc38a1*^+/+^ mice. Green marks «test start», whilst red marks «test end». (**B**) *Slc38a1*^-/-^ mice travel significantly shorter distances compared to *Slc38a1*^+/+^ mice in the elevated plus maze. (**C-E**) All three genotypes of *Slc38a1* have significantly more entries to the center zone, but the number of entries to the open and closed arms are similar. (**F-H**) All three genotypes spend significantly more time in the arms compared to the center zone. *Slc38a1*^-/-^ and *Slc38a1*^+/+^ spend significantly more time in the closed arms compared to open arms. The bars represent mean±SD. Dark grey shades represent *Slc38a1*^+/+^, grey *Slc38a1*^+/-^ and red *Slc38a1*^-/-^. Results for open arm, center zone and closed arms are shown by solid, vertical- and horizontal lines, respectively. Asterisks indicate the level of significance by one-way ANOVA with post hoc Tukey’s test (*p < 0.05, **p < 0.01, ***p < 0.001, ****p < 0.0001). The results are from a 5-minute test trial and included 17 *Slc38a1*^+/+^, 18 *Slc38a1*^+/-^ and 14 *Slc38a1*^-/-^ mice.

The significantly lower total distance travelled by *Slc38a1*^-/-^ mice in the elevated plus maze could be due to cognitive impairment, anxiety, depression and/or neuromotor dysfunction. Anxiety was therefore further examined by the open field test. Locomotor activity in the open field maze provides a good measure for anxiety-driven movements^32^. Anxious mice have less movements and prefer peripheral areas where they are less exposed compared to relaxed mice^33^. We performed the open field test to investigate spontaneous locomotor activity and to reveal any subtle evidence for anxiety-like behavior or lack of interest/motivation to explore the environment. *Slc38a1*^-/-^ spend significantly less time in the center area and significantly more in the peripheral area compared to *Slc38a1*^+/+^ mice (p<0.01; Fig. 4A, B), indicating reduced explorative behavior or more anxiety-like behavior. Consistent with this, the *Slc38a1*^-/-^ mice also rested significantly more in the peripheral area compared to the *Slc38a1*^+/+^ mice (p<0.01; Fig. 4C, D). Finally, grooming behavior (stereotypic episodes) in the center or peripheral areas are significantly reduced in *Slc38a1*^-/-^ mice compared to *Slc38a1*^+/+^ mice (p<0.01 and p<0.05, respectively; Fig. 4E, F).

**Figure 4.**
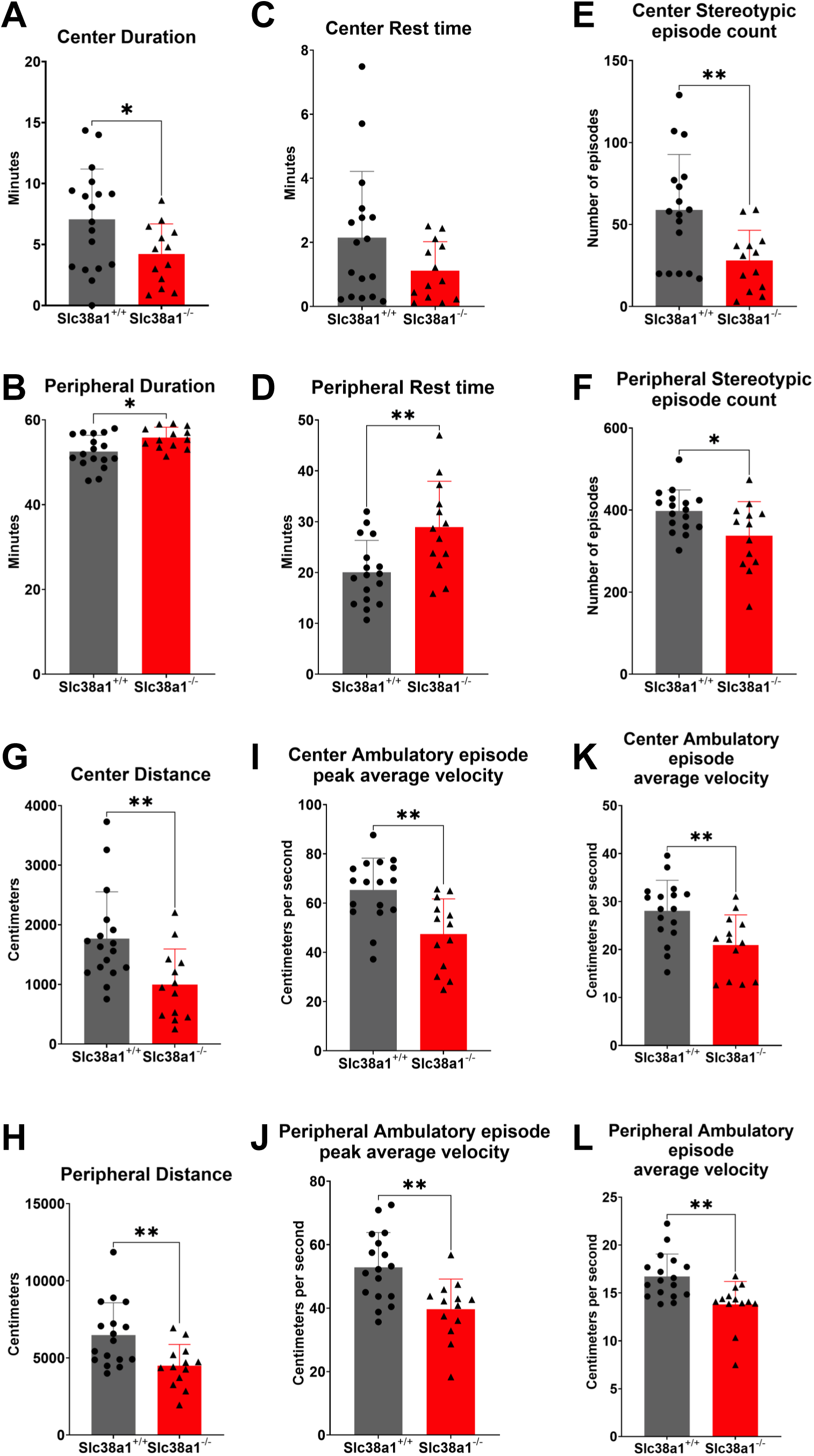
Genetic inactivation of *Slc38a1* in mice affects the duration, rest time, and stereotypic episode counts in the open field maze. The Open Field maze were assessed over 60 minutes to investigate any particular differences in locomotor activity. (**A, B**) *Slc38a1*^-/-^ mice spend significantly less time in the center region compared to *Slc38a1*^+/+^ mice. In the peripheral region, *Slc38a1*^-/-^ mice spend significantly more time than *Slc38a1*^+/+^ mice. (**C, D**) The parameter rest time indicates the length of time that the subject spent at rest. A resting period is defined as a period of inactivity greater than or equal to 1 second. *Slc38a1*^-/-^mice rest significantly more in the peripheral zone than *Slc38a1*^+/+^ mice. In the center zone, *Slc38a1*^-/-^ mice rest insignificantly less than *Slc38a1*^+/+^ mice. (**E, F**) Stereotypic episode count corresponds to the number of times that stereotypic behavior was observed in the animal. A one second or longer break in stereotypy is required to separate one stereotypic episode from the next. If the animal breaks the same beam (or set of beams) repeatedly, then the monitor considers that the animal is exhibiting stereotypy. This typically happens during grooming. *Slc38a1*^-/-^ mice exhibit significantly less grooming behavior in the center zone compared to *Slc38a1*^+/+^ mice. The peripheral stereotypic episode count is also significantly lower for *Slc38a1*^-/-^ mice compared to *Slc38a1*^+/-^. (**G, H**) Total distance in centimeters in the center zone and peripheral zone were analyzed to visualize any differences in ambulation. The distance acquired by *Slc38a1*^-/-^ mice is significantly lower in both center and in peripheral distance compared to *Slc38a1*^+/+^ mice. (**I, J**) Ambulatory episode peak average velocity is the highest value of the mean velocities for each ambulatory episode. There is significant reduction in *Slc38a1*^-/-^ mice compared to *Slc38a1*^+/+^ mice both in the centre and the peripheral ambulatory episode peak average velocity. (**K, L**) Ambulatory episode average velocity is the mean of the average velocities for each ambulatory episode. The ambulatory episode average velocity is significantly reduced, both in center and in the peripheral zones, in *Slc38a1*^-/-^ mice compared to *Slc38a1*^+/+^ mice. Bars represent mean±SD for *Slc38a1*^+/+^ (dark grey), *Slc38a1*^+/-^ (grey) and *Slc38a1*^+/+^ (red). Asterisks indicate level of significance by one-way ANOVA with post hoc Tukey’s test (*p < 0.05, **p < 0.01). n = 17 *Slc38a1*^+/+^, 18 *Slc38a1*^+/-^ and 13 *Slc38a1*^-/-^ mice.

The distance travelled by mice of the two genotypes shows significant reduction for *Slc38a1*^-/-^ mice compared to *Slc38a1*^+/+^ mice in both center and periphery (p<0.01; Fig. 4G, H). Furthermore, the ambulatory episode peak average velocity and average velocity was significantly reduced for *Slc38a1*^-/-^ mice compared to *Slc38a1*^+/+^ mice in the center as well as peripheral areas (Fig. 4I-L). Finally, stratifying the mice according to sex did not reveal any significant difference in center or peripheral duration between males and females in any of the three *Slc38a1* genotypes (Fig. S3).

Altogether, open field test data supports the role of *Slc38a1* in anxiety-like behavior. The significantly reduced total distance travelled in the elevated plus maze and in open field maze is also consistent with anxiety and/or depression.

### *Slc38a1* disruption does not affect sociability or preference for social novelty

We investigated a role for *Slc38a1* in social behavior by three-chambered sociability test. Both *Slc38a1*^-/-^ and *Slc38a1*^+/+^ mice explored the maze and had a similar number of entries to the left and right chambers, while the entry number to the middle chamber was significantly lower in *Slc38a1*^-/-^ compared to *Slc38a1*^+/+^ mice (Fig. 5A, B). When it comes to preferred chambers to spend time in, both *Slc38a1*^-/-^ and *Slc38a1*^+/+^ mice spent significantly more time in the chamber with stranger 1 than in the two empty chambers (Fig. 5C), with no difference between the genotypes. We next tested the preference for social novelty by introducing stranger 2 in the right chamber: Both *Slc38a1*^-/-^ and *Slc38a1*^+/+^ mice showed a significant preference for the chambers that contained a stranger mouse (Fig. 5D). Entry counts showed no difference between the two genotypes in the number of entries to the three chambers. Furthermore, both genotypes split their time equally between the chambers with stranger 1 and stranger 2 and spent significantly more time in these chambers than in the empty middle chamber (Fig. 5E-F).

**Figure 5.**
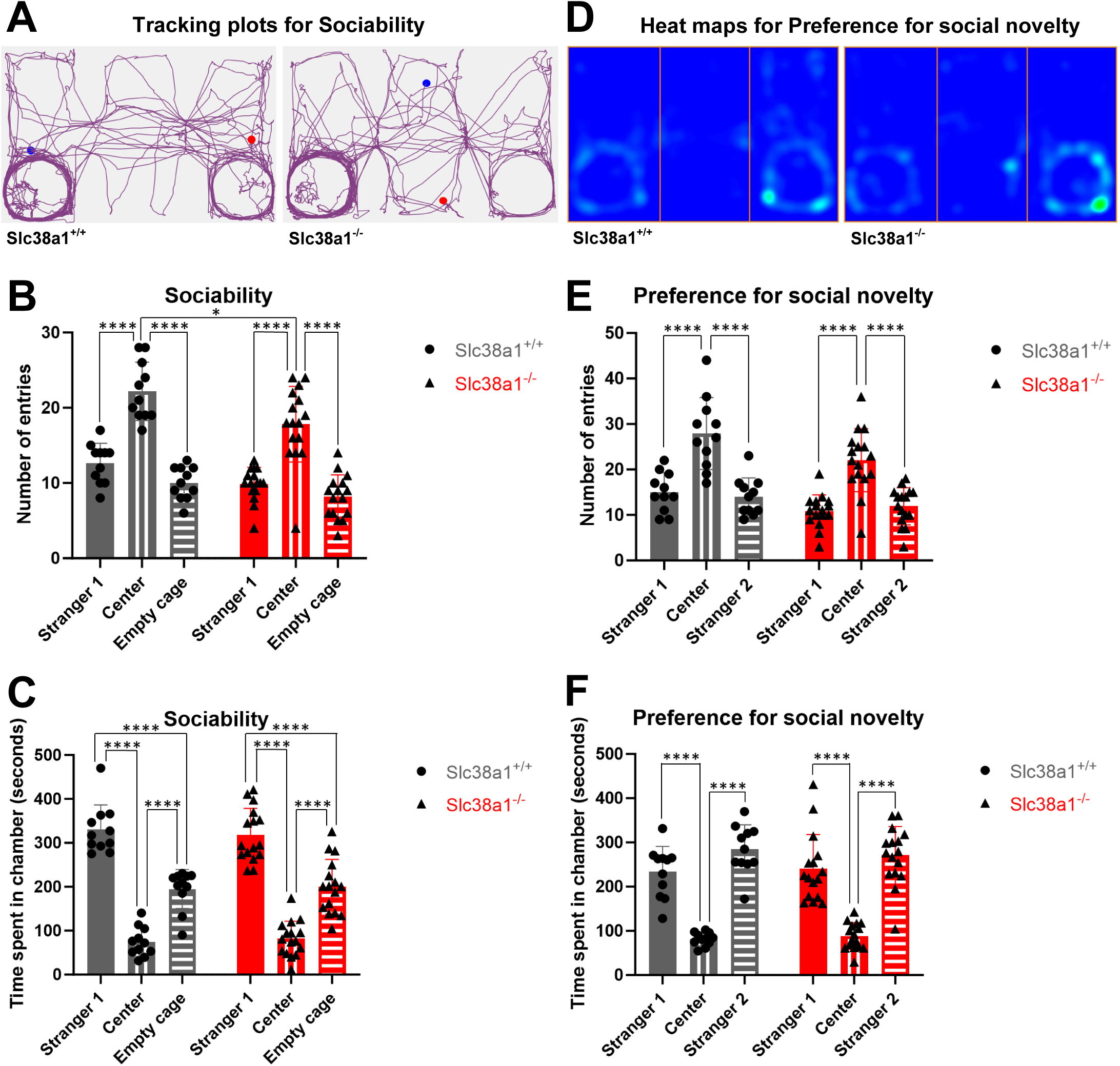
*Slc38a1*^+/+^ and *Slc38a1*^-/-^ mice show no differences in sociability or social novelty. Sociability and social novelty were tested in a three-chambered box with openings allowing passage between the three chambers. After habituation to the empty box, a mouse (stranger 1) is introduced under a cup in the left chamber for the “sociability test” while a cup in the right chamber remained empty. Later, a second mouse (stranger 2) was introduced under the cup in the right chamber for the “social novelty test”. (**A**) Representative tracing plots for two mice (one of each genotype) suggest that *Slc38a1*^+/+^ and *Slc38a1*^-/-^ mice display similar sociability. (**B**) *Slc38a1*^+/+^ and *Slc38a1*^-/-^ mice enter the middle chamber more frequently than the left chamber with stranger 1 or the empty chamber on the right. *Slc38a1*^+/+^ mice have a significantly higher number of entries in the middle chamber compared to *Slc38a1*^-/-^ mice. (**C**) Both *Slc38a1*^+/+^ and *Slc38a1*^-/-^ mice display sociability defined as spending more time in the chamber with stranger 1 than in the chamber with the empty wired cage or the center region. (**D**) Representative heat maps reveal that *Slc38a1*^+/+^ and *Slc38a1*^-/-^ mice are similar towards social novelty. (**E-F**) Upon introduction of stranger 2 in the right chamber, neither *Slc38a1*^+/+^ nor *Slc38a1*^-/-^ mice display preference for social novelty, i.e., both genotypes have similar number of entries and time spent in chamber 1 as well as in chamber 2. The bars show mean±SD. Bars with dark grey shades show data for the *Slc38a1*^+/+^ mice in the three chambers, while the bars in red shades represent data for the *Slc38a1*^-/-^ mice. Asterisks indicate level of significance by two-way ANOVA with post hoc Tukey’s test (*p < 0.05, ****p < 0.0001). 11 *Slc38a1*^+/+^ and 16 *Slc38a1*^-/-^ mice contributed to these experiments.

### *Slc38a1*^-/-^ mice do not expose motor impairments or metabolic changes

We tested whether *Slc38a1*^-/-^ mice were suffering from neuromuscular dysfunctions that could partly explain the observed reduction in locomotor activity. Fine motor coordination was assessed in the notch beam test where the mice were trained to walk across an elevated and narrow notched beam to a safe platform for three consecutive days and tested on the fourth day^26^. Less than two paw slips is considered normal for mice. We found no differences between *Slc38a1*^-/-^ and *Slc38a1*^+/+^ mice in latency to transverse the beam. None of the mice showed more than two paw slips and the number of paw slips did not differ between the genotypes (Fig. S4A, B).

To test for the potential impact of deletion of *Slc38a1* on metabolism or mobility, which subsequently could affect behavior, *Slc38a1*^+/+^, *Slc38a1*^+/-^ and *Slc38a1*^-/-^ mice were subjected to metabolic phenotyping in their natural home cages. Except for a significant difference in oxygen consumption between Slc38a1^+/+^ and Slc38a1^+/-^ males in the light phase, we found no differences in oxygen consumption, carbon dioxide production, or respiratory exchange ratio between the three genotypes, neither in the dark nor in the light phases (Fig. 6A-F). As expected, the activity of the mice was significantly higher during the dark phase compared to the light phase. However, no differences were detected between the three genotypes (Fig. 6G-H). Metabolic phenotyping also revealed no sex differences (Fig. 6A-H).

**Figure 6.**
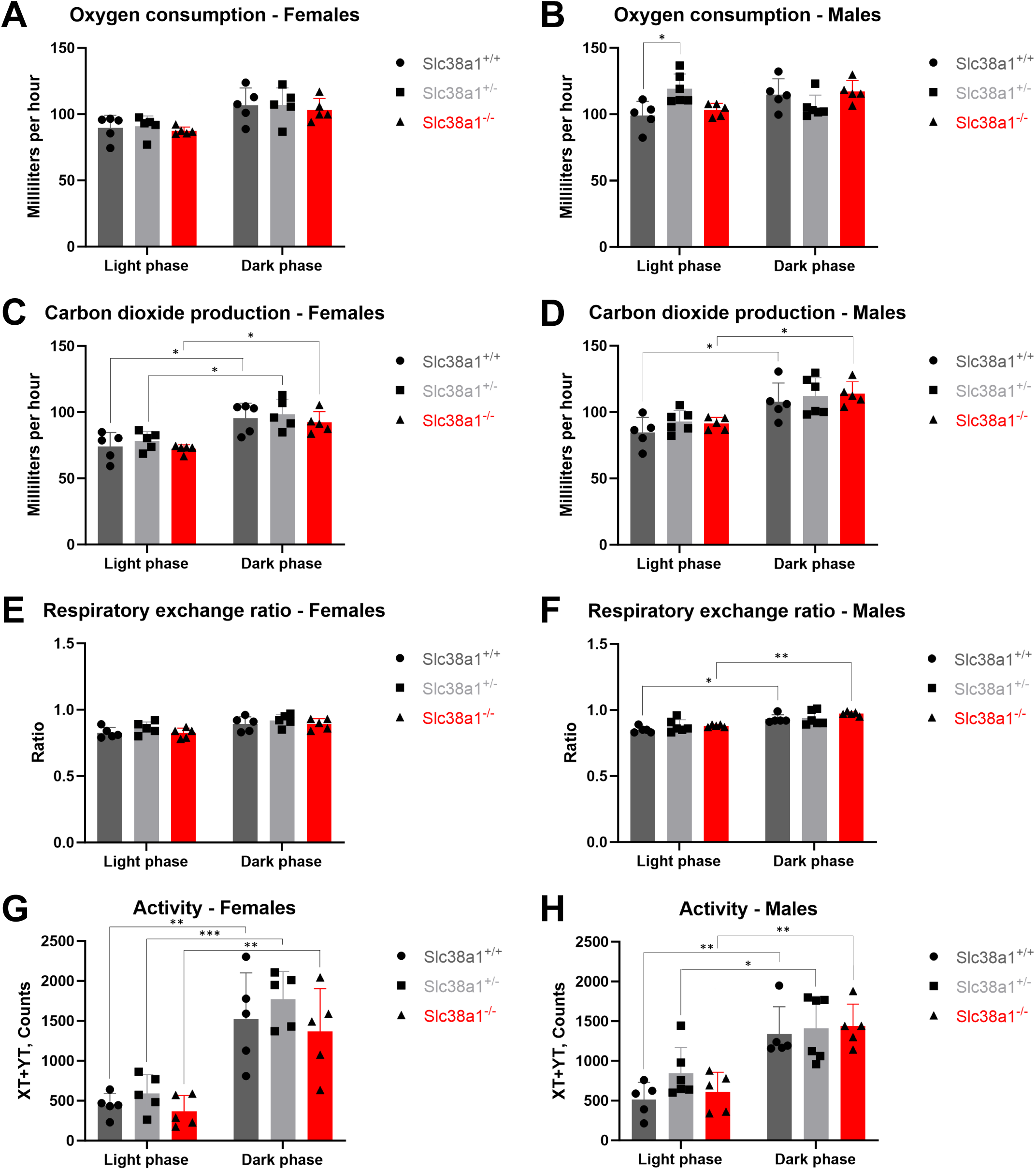
*Slc38a1* deletion does not impair metabolism in mice. Oxygen consumption, carbon dioxide production, and physical activity (movement in the XY-plane) were measured continuously for 48 hours. (**A**) The oxygen consumption, measured in milliliters per hour, show no differences between genotypes (female) in neither the light or dark phase. (**B**) Male *Slc38a1*^+/-^ mice show significantly higher oxygen consumption compared to *Slc38a1*^+/+^ males in the light phase. This difference is absent in the dark phase. (**C**) The carbon dioxide production, measured as milliliters per hour, in females differs significantly between the light and dark phases in all genotypes. However, there are no intergenotypic differences. (**D**) The carbon dioxide production in males differs significantly between the light and dark phases in *Slc38a1*^+/+^ and *Slc38a1*^-/-^, but not for *Slc38a1*^+/-^. There are no intergenotypic differences. (**E, F**) The respiratory exchange ratio, measured as ratio, shows no differences between genotypes or light/dark phases in females, but for males there are significant differences of respiratory exchange ratio between light and dark phases for *Slc38a1*^+/+^ and *Slc38a1*^-/-^, but not for *Slc38a1*^+/-^. There are no differences between genotypes. (**G, H**) As expected, the movement activity differs between the light and dark phase for all genotypes with higher activity rate during the dark phase. No significant changes are detected among the three genotypes. Metabolic phenotyping was done on male *Slc38a1*^+/+^ (n=5), *Slc38a1*^+/-^ (n=6), and *Slc38a1*^-/-^ (n=5) mice and female *Slc38a1*^+/+^ (n=5), *Slc38a1*^+/-^ (n=5), and *Slc38a1*^-/-^(n=5) mice. Data shown is based on mean values in the light (07am-07pm) and dark (07pm-07am) phases for the period. Respiratory exchange rate was calculated based on measured O_2_ and CO_2_. Bars show mean±SD.

## DISCUSSION

### Slc38a1 sustains spatial learning and memory

The lack of improvement in the latency to escape to the platform in the Morris water maze over five consecutive learning days combined with the significantly poorer performance of *Slc38a1*^-/-^ mice compared to *Slc38a1*^+/+^ mice in the probe trial on day six verify impaired spatial memory and learning when Slc38a1-mediated glutamine transport into GABA interneurons is blocked. Morris water maze has been considered a test for NMDA receptor-dependent long-term potentiation (LTP)^34,35^. We demonstrate that deletion of *Slc38a1* impairs spatial learning and memory in the Morris water maze despite the fact that *Slc38a1*^-/-^ mice have normal excitatory synaptic transmission, post-synaptic excitability, paired-pulse facilitation and LTP in the CA3-to-CA1^23^. Rather, our findings imply that Slc38a1-dependent glutamine transport into PV^+^- and SST^+^-interneurons and the subsequent GABA signaling make an important foundation for spatial learning and memory. This is consistent with NMDA-independent and/or GABA-induced LTP underpinning learning and memory^36,37^. Our data also harmonize with the reported correlation between GABA levels and/or the number of GABAergic interneurons on the one hand and learning and memory on the other; such a correlation has been suggested through several lines of evidence: 1) Attenuated GABA release by microwave exposure disables spatial memory in mice^38^, 2) Overexpression of GABA transporter 1 (GAT1) in the plasma membrane, which would be expected to facilitate reuptake of GABA from the synaptic cleft and thereby reduce GABA concentrations, has been reported to cause decline in associative learning capacity and decreased memory retention^39^, and 3) A reduction in PV^+^- or SST^+^-interneuron number and/or function would reduce GABA levels and deteriorate γ-oscillations. As expected, a selective loss of GABAergic neurons in rodents has been reported to result in poorer spatial learning and memory^40^. Further supporting a role for the GABA system in spatial learning and memory, there are also reports confirming a link between increased GABA signaling and enhanced memory formation: For instance, mice treated with the antiepileptic drug gabapentin, a GABA analogue activating GABA_B_ receptors, exhibited enhanced learning and memory in the Morris water maze: gabapentin-treated mice spent significantly more time in the target quadrant, and the swimming distance to the platform was decreased compared to untreated mice^41^. Increasing endogenous GABA levels by administration of γ-vinyl-GABA, an inhibitor of GABA catabolism, significantly and dose-dependently reduces escape latency^42,43^. Finally, inhibition of GAT1 by, e.g., NNC-711, is expected to elevate extracellular levels of GABA. Both mature and aged rats treated with NNC-711 showed reduced escape latency in the Morris water maze and reduced amnesia in a passive avoidance task^44^. Altogether, our data demonstrate a role for Slc38a1-dependent GABA neurotransmission in spatial learning and memory. The shorter swimming distance and slower speed together with significantly less time spent in North and platform zones by *Slc38a1*^-/-^ mice and no involvement of motoric and/or motivational deficits support that these mice experience cognitive impairment^42,43^.

### Disruption of *Slc38a1* induces depressive-like behavior

The forced swim test revealed depressive-like behavior in *Slc38a1*^-/-^ mice compared to *Slc38a1*^+/+^ mice. This is compatible with numerous reports highlighting dysfunctional GABA signaling in depression. Significantly reduced GABA levels are detected in the plasma, the cerebrospinal fluid (CSF), and the cerebral neocortex of individuals diagnosed with depressive disorder^45,46,5^. Magnetic resonance spectroscopy studies also show diminished GABA levels in the brain cortices of depressed patients^47^. Selective reduction of GAD67 in humans or GAD67^+/GFP^ transgenic mice is also associated with depression^48,49^. In line with this, alterations in the number or affinity of GABAA-receptors have been shown in MDD^50,51^.

Deteriorated SST^+^-interneuron function is another hallmark of depression. *Post-mortem* studies as well as investigation of patients, provide a strong association between decreased levels of SST/calbindin and MDD^52,53,5^. Most of these and other studies also testify that MDD, together with numerous other behaviors related to mood and emotions, affect particularly the prefrontal cortex (PFC), hippocampus, anterior cingulate cortex (ACC), amygdala and/or subcortical reward circuits^54,55,56^. Slc38a1 is enriched in PV^+^- and SST^+^-interneurons^19,20,21^ ‒ with impact on neurotransmitter GABA synthesis, synaptic plasticity and the excitatory/inhibitory balance^23,22^ ‒ and we now demonstrate an association with depression. Thus, this together with our current data bolsters the GABAergic deficit theory for MDD^3^, and suggests that Slc38a1 may be a central protein in the pathophysiology of depression. Furthermore, MDD is commonly associated with neuropsychiatric impairments^57,58^, and we now demonstrate impaired spatial learning and memory along with depression-like behavior in the *Slc38a1*^-/-^ mice.

### *Slc38a1*^-/-^ mice may harbor latent anxiety-like behavior

In the open field test, we observed a significant reduction in center duration, increased peripheral duration and rest time together with reduced center and peripheral stereotypic episodes in *Slc38a1*^-/-^. The *Slc38a1*^-/-^ mice also travelled significantly less compared to *Slc38a1*^+/+^ mice in the elevated plus maze and open field. This cannot be due to locomotor disabilities since the notch beam test and metabolic phenotyping did not differentiate between the two genotypes. Our data from the open field test indicates waning interest in exploration over time and is compatible with depressive-like behavior and/or the presence of latent anxiety.

Depression and anxiety, do often occur together, and a potential role of Slc38a1 in anxiety would not be an unexpected finding as anxiety is associated with GABAergic dysfunction and GABA is required for both acquisition, consolidation, reconsolidation, and extinction of fear memory^59,60^. Furthermore, anxiety and MDD show comorbidity and respond to the same treatments^61;62,63^. Enhancing GABA transmission by drugs, such as benzodiazepines, tiagabine, and neurosteroids, is anxiolytic^64,65^.

Also, in regards to brain regions, anxiety-like behavior is associated with several of the same brain regions as depression-like behavior, including PFC, hippocampus, ACC and particularly inhibitory networks in the amygdala^65,66,67,68^,; regions where PV^+^-interneuron associated γ-oscillatory activity is associated with anxiety^68^ and in which Slc38a1 is highly expressed^19,20,23^. Stimulating GABAergic neurotransmission by infusion of GABA or GABA receptor agonists, e.g., in the amygdala, reduced fear and anxiety while infusions of GABA antagonists tend to have anxiogenic effects^69^. Also, 50% of the interneurons residing in the basolateral amygdala are PV^+^ associated with γ-oscillations which synchronize the excitatory neurons, regulate their output and generate and maintain oscillatory activity facilitating information processing^70,71^. Impairment of this PV^+^-interneuron associated γ-oscillatory activity is associated with anxiety^68^. Interpreted in light of the literature mentioned above, our data support that aberrant GABA signaling and/or γ-oscillations associated with Slc38a1 glutamine transport into PV^+^- and SST^+^-interneurons may contribute to the development of anxiety. Moreover, GABAergic signaling boosts dendrite outgrowth and development of dendritic maturation and arborization^72,73^, and we have previously demonstrated that *Slc38a1*^-/-^ mice have perturbed maturation of dendrites, including shortened dendritic branch length and reduced complexity of dendritic arbor^21^. Indeed, defects in dendrites, spines and/or synaptogenesis are associated with depression and anxiety^51,74^, and a pathologic variant of *Slc38a1* has been reported in a family with depression and suicidal behavior^75^.

### Slc38a1 impairment do not cause schizophrenia-like or ASD-like behaviour

Slc38a1 impairment had no impact on sociability or social novelty in the three-chambered box. These findings show that *Slc38a1*^-/-^ mice do not display locomotor hyperactivity, stereotypic movements or social deficits which are key attributes of schizophrenia and ASD^76,77,49,10^. This is surprising since declined GABA levels can induce and/or exacerbate symptoms of schizophrenia^78,79,17,80^. Likewise, abnormal GAD67 and/or PV^+^-interneuron activity are strongly associated with schizophrenia^8,81,79,82^. Genetic knock-down of *GAD67* in PV^+^-interneurons, e.g., induces schizophrenia-like behavior in mice^83,84^. In addition, GABA neurotransmission, GAD67 activity and PV^+^-interneurons are critical for γ-oscillations, which are important for optimizing cognitive functions, such as working memory, attention, and consciousness^85,12,15,81,,86,84^. Indeed, optogenetic activation of PV^+^-cells has been shown to be sufficient for inducing γ-rhythms and controlling sensory responses to enhance cortical circuit performance^87^. In schizophrenia, abnormal oscillations, particularly in the dorsolateral PFC, impair cognitive functions and promote hallucinations^78,88^. Such abnormal γ-oscillations are also important for the interaction and transfer of information between different brain regions, which is a hallmark of schizophrenia^12^. ASD has some of the same clinical features and pathogenic mechanisms as schizophrenia^77,11^. Also, in ASD there is dysfunctional GABAergic signaling, and γ-oscillations are disrupted^89,90^. Moreover, the fragile X messenger ribonucleoprotein (FMR1) knock-out mouse model of ASD displays aberrant γ-oscillations, malfunction of PV^+^-interneuron, and imbalance in excitatory/inhibitory signaling^91,87,10^. Altogether, our data show that the reduced total levels of GABA and significantly slower oscillations in the γ frequency band in Slc38a1^-/-^ mice^23^ seem not to be enough to perturb sociability, social novelty, or induce other symptoms of schizophrenia and/or ASD, suggesting a slightly different pathogenesis or requiring different quality of the γ-oscillations.

### Proteins involved in GABA signaling and γ-oscillations may differentially shape multiple neuropsychiatric disorders

A prevailing hypothesis suggests that the ultimate action of depression therapy is to rectify GABA deficit and/or to upregulate GABA systems^17,92^. This is seen, e.g., after selective serotonin reuptake inhibitors (SSRIs), electroconvulsive therapy and transcranial magnetic stimulation (TMS)^93,3,94^. In accord with this, treatment with the GABAA receptor agonist diazepam, which certainly has antidepressive effects^95,96^, upregulates Slc38a1. Increasing expression and/or activity of Slc38a1 increases GABA levels and GABAergic signaling and has trophic effects required for normal brain development^97,21^.

However, the GABAergic system consists of several pathways which are differentially regulated and with the potential for differential outcomes. Disruption of glutamic acid decarboxylase 65 (GAD65) reduced GABA levels in the amygdala and exposed anxiety and pathologic fear memory but reduced immobility ties in the forced swim test, i.e., antidepressant behavior^98,99^. The situation is partly opposite in our study: *Slc38a1*^-/-^ mice spend more time immobile in the forced swim test than their *Slc38a1*^+/+^ littermates, i.e., more depressive-like behavior, and reveal anxiety in the open field maze, but not in the Plus maze. Despite both genotypes having GABA deficit, the different phenotypes are due to dissimilar pathophysiology: *Slc38a1* disruption does not diminish GAD65 and compensates loss of GABA release by increasing GAD67 significantly and maintains basic GABA release at the nerve terminals^23^. Thus, although both anxiety and MDD are associated with GABA deficit, they may be associated with different metabolic pathways and/or pools of GABA, differentially impacting tonic and/or phasic GABA neurotransmission and involving different neuronal circuits. On top of that, γ-oscillations and impact on membrane trafficking and function of GABA receptor units, may shape the neuropsychiatric diseases. This may explain why disruption of Slc38a1 did not reveal behavior compatible with schizophrenia or ASD even though transplantation of interneuron precursor cells in the adult hippocampus has previously been reported to be successful in reversing some psychosis-relevant features in a mouse model of hippocampal disinhibition^100^.

