## Supplementary information for "Disruption of glutamine carrier *Slc38a1* causes cognitive impairment, anxiety and depressive-like behavior"

**Abbreviated title: Neuropsychiatric aspects of *Slc38a1***

### MATERIAL AND METHODS

#### Animals

This study was conducted in strict accordance with the recommendations described in the Norwegian Animal Welfare Act and the European Union Directive on the Protection of Animals used for Scientific Purposes (2010/63/EU). All experiments have been approved by the Norwegian Food Safety Authority (FOTS application numbers 9200, 10902, 21009 and 30120). Mice were housed in a room with fixed 12-hour light/dark cycle, temperature set to 22 ± 1 °C with relative humidity at 50 ± 10%. The animals were stalled in groups up to 9 mice in Makrolon GreenLine cages (Sealsafe Plus GM500 or GM900, Buguggiate, Italy). The cages were always ventilated with 100% fresh air except during experiments. The cages were enriched with paper for nest-building, a wooden stick, a paper roll or a small plastic house and a running wheel. Autoclaved food (RM3 from Special Diets Service (UK)) and water were provided *ad libitum*.

The molecular impact of *Slc38a1*$^{-/-}$ mouse model used in these studies has previously been rigorously characterized[1,2]. Animals used for the study were backcrossed in to C57BL/6J for 10 generations. Mice that were wild type (*Slc38a1*$^{+/+}$), heterozygous for the mutation (*Slc38a1*$^{+/-}$), or genetically inactivated for *Slc38a1* (*Slc38a1*$^{-/-}$) were all originating from the same breeding colony. The mice were earmarked for identification at 4 weeks of age. The biopsies taken during earmarking were used for genotyping. Animals included in the study were of both sexes and aged 10-15 weeks, except for metabolic phenotyping where they were 22-23 weeks. All behavioral tests were performed in the same room, at the same time of the day (between 1:00 p.m. and 6:00 p.m.) and by the same experimenter to minimize any effects of environmental factors. Before each behavioral test all animals had a habituation period of one hour to acclimate to the experimental environment.

#### Morris water maze

The Morris water maze setup consisted of a white, circular pool with a diameter of 150 cm and depth of 30 cm. The pool was placed in a room with white walls with high-contrast visual cues on each wall, white floors, and ceiling. The visual cues, with different colors and dimensions, were kept in the exact same place throughout the whole experiment. The pool was filled to a depth of 21,5 cm with water which was made opaque by adding a small amount of non-toxic Jotun SENS paint (Jotun, Norway), the water temperature was maintained at 25 ± 1 °C, and the light was set to approximately 40 lux. The Morris water Maze arena was digitally divided into four quadrants (north, south, east and west) by an advanced video tracking system (ANY-maze; Stoelting Europe, Ireland). A white platform with a diameter of 11 cm was then placed 29,5 cm inwards from the pool edge in the northern quadrant and submerged 0,5 cm below the water surface. After the last trial each day, the wet animals were dried in a paper towel and placed in a clean cage, heated with an infrared heating lamp, for at least 5 minutes, before being returned to their home cages. The pool was emptied each day and refilled with opaque water the next day. In addition, the apparatus was dried with paper towels and cleaned with 70% ethanol at the end of each trial day.

Since behavioral experiments have not been conducted on *Slc38a1* mice previously, two protocols lasting for six or ten days were tested. The10-day protocol did not provide additional significant advantages compared to the 6-day protocol. Therefore, the tests were performed using the 6-day protocol. The platform was placed in the same position throughout the learning trials (day 1-5) but was removed from the pool during the probe test (day 6). The pool was placed in a separate room from the experimenter.

Test animals were trained to learn the position of the platform by identifying distal visual cues and using these in order to escape the water. Each animal was given in total 20 learning trials: four trials/day for five consecutive days. Animals were lifted by the base of their tail and gently released into the water with their head positions towards the edge of the pool. Then, the experimenter quickly left the testing area. For each learning day, all animals were released at four different locations (west, east, southwest and southeast). The order of locations was randomized for each day but was the same for all animals. The animals were then given 60 seconds to swim and search for the platform. If an animal could not locate the platform within the 60 second trial, it was placed on the platform for 30 seconds by the experimenter. However, if an animal located the platform, the trial was manually stopped by the experimenter, and the animal was allowed to stay on the platform for 30 seconds. To examine spatial reference memory, a probe test was performed on day 6, 24 hours after the last training session. For the probe test the platform was removed, and all animals were released in a new start position (south). The probe test consisted of only one trial, and animals were allowed to swim freely for 60 seconds. Animals floating for more than 30 seconds were excluded from the probe test.

ANY-maze automatically measured latency before entering each zone for the first time, number of entries in each zone as well as time spent in each zone. The different zones were marked as “north”, “south”, “east”, “west”, “platform” and “peri-platform” (defined as 3 cm wider than the platform zone). Repeated Measures ANOVA with post hoc Tukey’s test was performed to analyze both data from the trial days and the test day.

**Forced swim test**

During the forced swim test three animals were tested simultaneously, each animal was placed in its own cylinder made of transparent glass with measurements of 40 cm in height and 17 cm in diameter. The glass cylinders were filled with clear water up to 20 cm in height to prevent the mice from resting with their tails on the bottom of the cylinder during the experiment. The water temperature was 25 ± 1 °C and the light intensity above the cylinders was approximately 40 lux. Turquoise colored partitions were placed in between and behind the glass cylinders to ensure the animals could not see each other during the test. The test took place in a separate room from the experimenter. However, the experimenter was still able to observe the mice during the procedure to ensure that the test animals never appeared to be in serious distress (severe tiredness, not floating or swimming, etc.). A GoPro™ camera was placed in front of the apparatus, at the height of the water line, in a way that allowed clear observation of the behavior of all animals placed in the glass cylinders.

The video recording was started before placing the mice gently into the water tanks by holding the base of their tail. After a six-minute time lapse, the video recording was stopped, and animals were removed from the water by their tails in the same order that they were put in. Before returning to their home cages, mice were dried using paper towels and placed in cages heated by an infrared lamp. The lamp was placed so that only half of the cage was heated, and the mice could choose to stay in the heated or the non-heated area. The water was changed after every session to avoid any disturbance on behavior of the next animal[3].

Only the last four minutes of the forced swim test were analyzed due to the disturbances on measurements caused by excessive activity in most mice at the beginning of a procedure of this sort[4]. The experimenter used a stopwatch to measure mobility time, and the total amount of mobility time was subtracted from 240 seconds which was stated at the time the animal was immobile. The definition of mobility time used in the forced swim test is "*any movements other than those necessary to balance the body and keep the head above the water*"[4]. The experimenter was trained to correctly identify movements that are counted as mobility in forced swim test. In addition, the experimenter was blind to the genotype of the mice and recordings were analyzed twice to minimize bias. Data collected were analyzed by One-way ANOVA followed by post hoc Tukey's test to compare immobility time between *Slc38a1*$^{+/+}$, *Slc38a1*$^{+/-}$ and *Slc38a1*$^{-/-}$ mice. Two-way ANOVA with post hoc Tukey's test was used to compare immobility time between both genotype and gender.

**Elevated Plus Maze**

The elevated plus maze apparatus was shaped as a plus sign (+) and elevated 50 cm above the floor. Each arm of the plus maze was 25 cm long and 5 cm wide and had a white color. Two facing arms were enclosed by 16 cm high walls (closed arms), and the arms perpendicular to the closed arms had an edge which was only 0,5 cm (open arms). The apparatus was placed in a separate room from the experimenter and illuminated by indirect light which was maintained at 40 lux.

The animal was placed in the center zone of the maze with its head directed toward a closed arm. The animal was then allowed to explore the maze freely for five minutes. Each mouse got only one test trial. A video tracking camera was placed above the apparatus in a manner so the shadows from the walls did not cover the platform and obscure the recording of the animal's movements. The recording was connected to an advanced video tracking system (ANY-maze). This software automatically collected data, such as total distance, number of entries to open or closed arms or the center zone, as well as the accumulated time spent in open arms, closed arms, or the center zone. Before placing a new animal for testing in the maze, the apparatus was thoroughly cleaned with soap-water and dried with paper towels. After each test day, the apparatus was cleaned with 70% ethanol.

One-way ANOVA with Tukey's post hoc test was used to compare total distance/number of entries/time spent in the different areas of the apparatus between the three *Slc38a1* genotypes.

**Open field maze**

The open field tests were performed in a VersaMax Animal activity monitor (AccuScan Instruments, Inc., USA). The apparatus consists of a square-shaped white 40 cm x 40 cm arena, with 30 cm plexiglass walls. Hollow lids made of plexiglass covered the top of the apparatus to prevent the mice from jumping out. The illumination level was maintained at 40 lux, and the lightning was adjusted to not leave shadows or dark corners in the open field arena.

Eight mice were tested in separate arenas at the same time. After a 60-minute habituation period, the mice were placed in the centre of the maze and were allowed to explore the maze freely for 60 minutes. Mice movements were tracked for all 60 minutes by infrared sensors in the floor of the square activity monitor. The arena was digitally divided into a center zone (defined as a 20 x 20 cm area, 10 cm away from the walls), and a peripheral zone consisting of the rest of the arena. The smell of male mice can make female mice anxious; therefore, experiments were first performed on all female mice followed by the male mice. This applies for all experiments. In between tests, the apparatus was thoroughly cleaned with towels soaked with soap-water first, followed by 70% ethanol and completely dried with paper towels.

The distance travelled (both in the peripheral and the center area) and the speed of the movements were detected. All statistical analyses were performed by GraphPad Prism Version 9.0. Repeated Measures ANOVA with post hoc Tukey's test was used to compare different parameters between *Slc38a1*$^{+/+}$, *Slc38a1*$^{+/-}$ and *Slc38a1*$^{-/-}$ mice.

**The three-chambered social test**

The three-chambered sociability test setup was divided into three-chambers by removable partitions. The partitions had openings to allow the test mouse to move freely between the chambers. Each of the distal chambers contained a wired cage placed on a disk. A weighted cup was placed on top of each wired cage to prevent the test mouse from staying on top of these cages.

This experiment consists of three phases: a habituation period, the sociability test and a preference for social novelty test. During the habituation period the test mouse was placed in the middle chamber with the dividers open to allow the mouse to explore the apparatus for 10 minutes. Both wired cages were empty in this period. After the habituation period, the test mouse was guided back into the middle chamber, and the dividers were gently closed. For the sociability test, a C57Bl/6J mouse which was unfamiliar to the test mouse (stranger 1) was placed in one of the wired cages, while the one on the opposite side was still empty. The dividers were then raised, and the test mouse was allowed to freely explore the arena for 10 minutes. Immediately after the sociability test, the test mouse was confined to the middle chamber. A new stranger mouse (stranger 2) was placed in the previously empty wired cage, while stranger 1 remained in its wired cage. The two strangers came from different home cages, and there was no interaction between mice prior to the experiment. The sex and age of the stranger mice correlated with the sex and age of the test mouse. Prior to the experiment, the mice to be used as stranger mice were habituated to the wired cage for 10 minutes daily for three consecutive days. During the 10-minute daily habituation, the experimenter monitored the mice to ensure that the mice used as strangers acted in a consistent manner with minimal aggressive behavior[5; 6]. With both of the stranger mice in place, the dividers were again removed, and the test mouse was allowed to move freely for another 10 minutes to test for preference for social novelty.

The entire experiment was videotaped from above by a video tracking camera connected to the ANY-maze software which automatically measured the time spent in each chamber and the number of entries into each chamber. Preference for social novelty was defined as more time spend with stranger 2 than stranger 1[5,6]. This experiment was performed on a laboratory bench in a separate room from the experimenter to minimize gradients in light, sound, smell and other environmental conditions that could lead to a side preference. To minimize odor transfer, the apparatus, wired cages and disks were firmly cleaned with soap and water and then wiped with paper towels before each new test subject. After each test day, everything (the apparatus and all items the mice had touched) was cleaned with 70% ethanol and left to air dry.

Two-way ANOVA with post hoc Tukey’s test was used to compare number of entries/time spent in the chamber containing the stranger 1 mouse and number of entries/time spent in the chamber containing the empty wired cage/stranger 2 mouse.

**The notched beam test**

The notch beam test apparatus was a 1-meter-long beam consisting of 15 mm blocks with 15 mm gaps in between the blocks. A black box was placed at the end of the beam as the finish point. The apparatus was elevated 50 cm above the table. A nylon hammock was placed below the beam (10 cm above the table) to cushion any eventual falls. A GoPro™ camera was placed in front of the apparatus in a way that allowed clear recording of the center 50 cm of the length of the beam.

This experiment consisted of three training days, followed by a test day. On training days, the *Slc38a1*$^{+/+}$ and *Slc38a1*$^{-/-}$ mice were trained to cross the beam three times each, without stalling. If a mice stalled to sniff or look around without moving forward, the experimenter encouraged the mice to move forward by gently touching the tail or pushing the mouse from behind. When the mice reached the black box, they were allowed to rest there for 15-30 seconds before the next trial. After each session, the mice were returned directly to their home cages.

On the test day, the mice were videorecorded while crossing the beam. Time spent to transverse the center 50 cm of the beam was measured by a stopwatch, and the number of paw slips (defined as the foot coming off the top of the beam[7]) was counted by analyzing the video recordings. Two successful trials in which the mouse did not stall on the beam were averaged and further analyzed by unpaired t-test. The beam and the black box were cleaned with towels soaked with 70% ethanol and allowed to dry completely before the next mouse was tested.

**Metabolic phenotyping**

The average ages for the male and female groups at the time of analyses were 23.10 and 22.53 weeks, respectively. Male and female *Slc38a1*$^{+/+}$, *Slc38a1*$^{+/-}$ and *Slc38a1*$^{-/-}$ mice were placed in a metabolic cage system consisting of 20 individual cages (Phenomaster, TSE Systems, Germany) as described previously[8]. After 36-40 hours of single housing and acclimatization in the metabolic cages, indirect calorimetric measurement of oxygen consumption and carbon dioxide production were performed with the following settings: gas flow rate at 0.42 L/min, measurement time for 10 seconds, and gas exchange measurement in 20 minutes intervals. Physical activity was measured as movement in the XY-plane[8]. Data collected over a 48-hour period were used to calculate mean values in the light (12 hours) and the dark (12 hours) phases, respectively. Respiratory exchange ratio (RER) values were calculated based on measured $O_2$ and $CO_2$ values. Body weights were measured prior to metabolic phenotyping.

### SUPPLEMENTARY FIGURES

**Supplementary Figure S1. No gender differences observed among the *Slc38a1*$^{+/+}$, *Slc38a1*$^{+/-}$ and *Slc38a1*$^{-/-}$ mice in spatial learning and memory in the Morris Water Maze**

(**A-I**) Data from Morris water maze obtained in Figure 1 were separated and stratified for sex. There are no significant differences between males (M) and females (F) on latency to first entry, number of entries or time spent in the north zone, the peri-platform zone or the platform zone among the three genotypes investigated (*Slc38a1*$^{+/+}$, *Slc38a1*$^{+/-}$ or *Slc38a1*$^{-/-}$ mice).

Bars represent mean±SD for *Slc38a1*$^{+/+}$ (dark grey) *Slc38a1*$^{+/-}$ (grey) and *Slc38a1*$^{+/+}$ (red). n = 14 males and 12 females *Slc38a1*$^{+/+}$, 5 males and 8 females *Slc38a1*$^{+/-}$ and 11 males and 15 females *Slc38a1*$^{-/-}$ mice.

**Supplementary Figure S2. *Slc38a1*$^{-/-}$ mice have significantly less entries to all three zones and spend less time in the center zone in the elevated plus maze**

(**A-C**) *Slc38a1*$^{-/-}$ and *Slc38a1*$^{+/-}$ mice have a significant lower number of entries in all zones (open arms, closed arms and center) compared to *Slc38a1*$^{+/+}$ mice. (**D-F**) There is no significant difference in the time spent in neither open arms nor closed arms between *Slc38a1*$^{+/+}$, *Slc38a1*$^{+/-}$ and *Slc38a1*$^{-/-}$ mice. However, *Slc38a1*$^{+/-}$ and *Slc38a1*$^{-/-}$ mice spend significantly less time in the center of the elevated plus maze compared to *Slc38a1*$^{+/+}$ mice.

These data are obtained from 5 minutes test trials and include 17 *Slc38a1*$^{+/+}$, 18 *Slc38a1*$^{+/-}$ and 14 *Slc38a1*$^{-/-}$ mice. Asterisks indicate level of significance by one-way ANOVA with post hoc Tukey's test (*$p < 0.05$, **$p < 0.01$, ***$p < 0.001$, ****$p < 0.0001$).

**Supplementary Figure S3. Open field maze show no gender differences in the *Slc38a1*$^{+/+}$, *Slc38a1*$^{+/-}$ and *Slc38a1*$^{-/-}$ mice**

We stratified the genotypes in males and females and analyzed results obtained open field test. This test shows no significant differences in center or peripheral duration between males and females.

Bars represent mean±SD for *Slc38a1*$^{+/+}$ (dark grey), *Slc38a1*$^{+/-}$ (grey) and *Slc38a1*$^{+/+}$ (red). Solid bars represent males, while hatched bars represent females. n = 10 males and 7 females *Slc38a1*$^{+/+}$, 8 males and 10 females *Slc38a1*$^{+/-}$ and 5 males and 8 females *Slc38a1*$^{-/-}$ mice were included in open field tests.

**Supplementary Figure S4. *Slc38a1*$^{-/-}$ mice do not have motor imbalance or miscoordination**

*Slc38a1*$^{+/+}$ and *Slc38a1*$^{-/-}$ mice underwent notched beam test after three days of training. (**A**) Average times for crossing the notched beam for *Slc38a1*$^{+/+}$ and *Slc38a1*$^{-/-}$ mice did not deviate significantly (3.68±0.83 and 4.03±1.05 seconds±SD, respectively). (**B**) Both *Slc38a1*$^{+/+}$ and *Slc38a1*$^{-/-}$ mice had less than 0,5 paw slips on average and none of the test animals fell off the beam.

Bars show mean±SD. Unpaired t-test was used to analyze the data. 11 *Slc38a1*$^{+/+}$ and 13 *Slc38a1*$^{-/-}$ mice were tested by notched beam test.
