## Supplementary figures for "Disruption of glutamine carrier *Slc38a1* causes cognitive impairment, anxiety and depressive-like behavior"

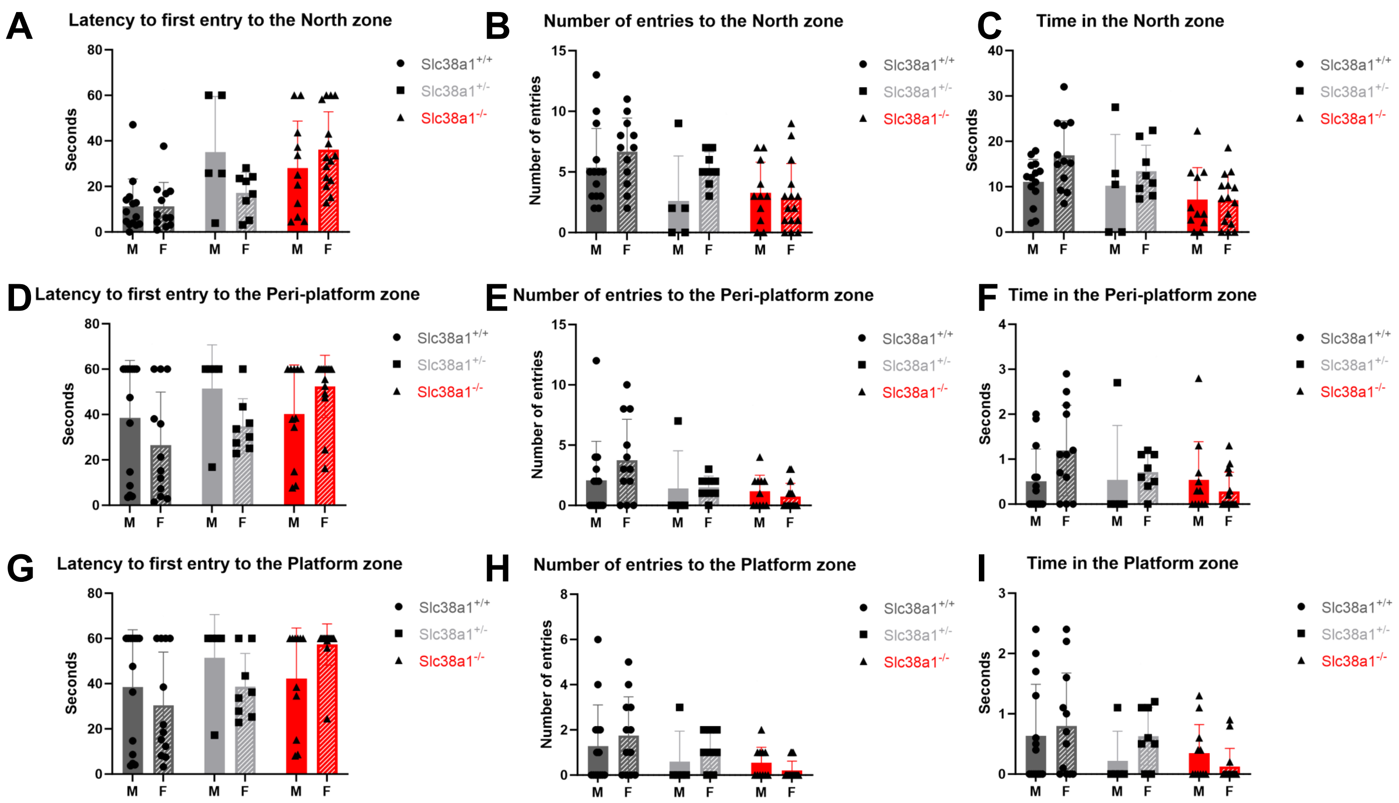


**Supplementary Figure S1. No gender differences seen among the Slc38a1$^{+/+}$, Slc38a1$^{+/-}$ and Slc38a1$^{-/-}$ mice in spatial learning and memory in the Morris Water Maze (MWM)**

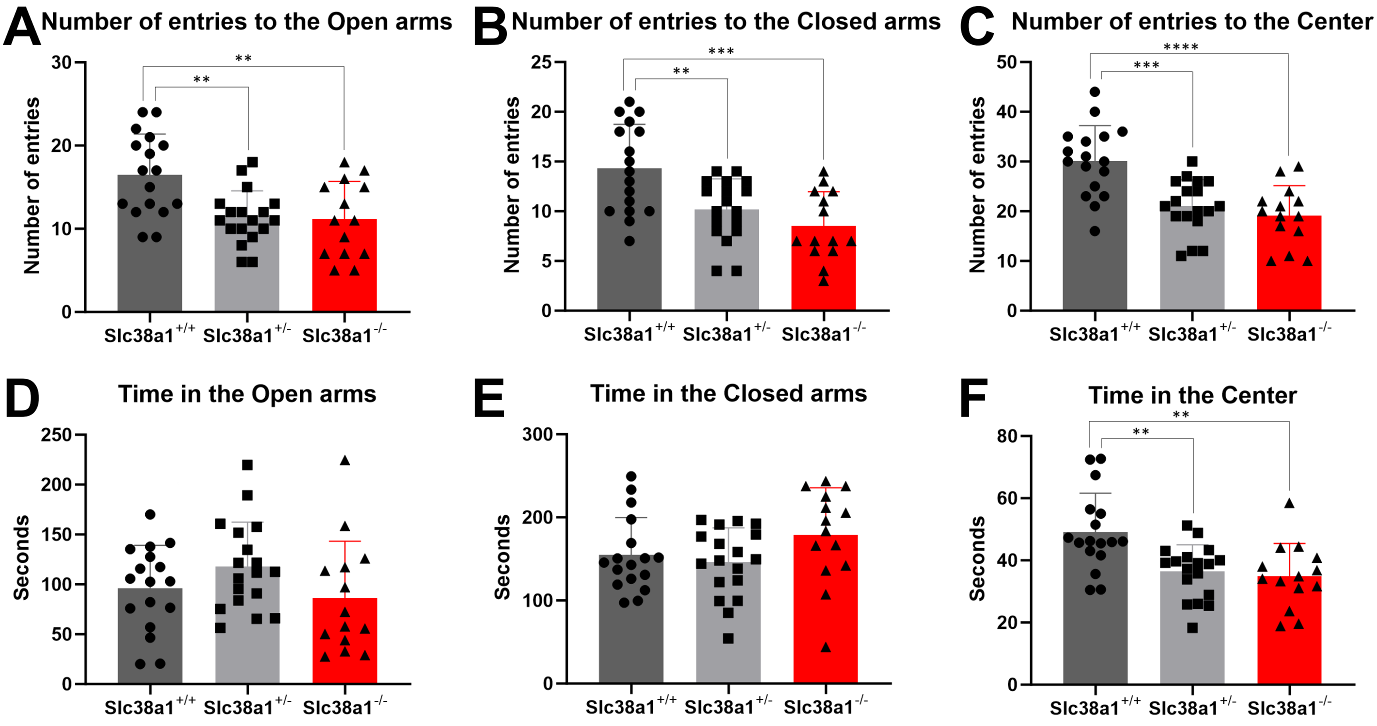


**Supplementary Figure S2. Slc38a1$^{-/-}$ mice have significantly less entries to all three zones and spend less time in the center zone in the elevated plus maze (EPM)**

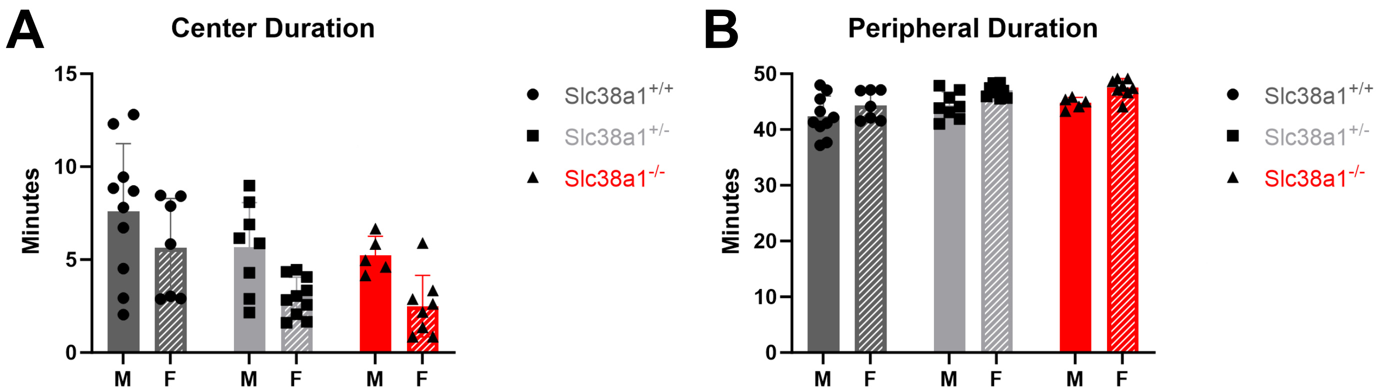


**Supplementary Figure S3. Open field (OF) test show no gender differences in the Slc38a1$^{+/+}$, Slc38a1$^{+/-}$ and Slc38a1$^{-/-}$ mice**

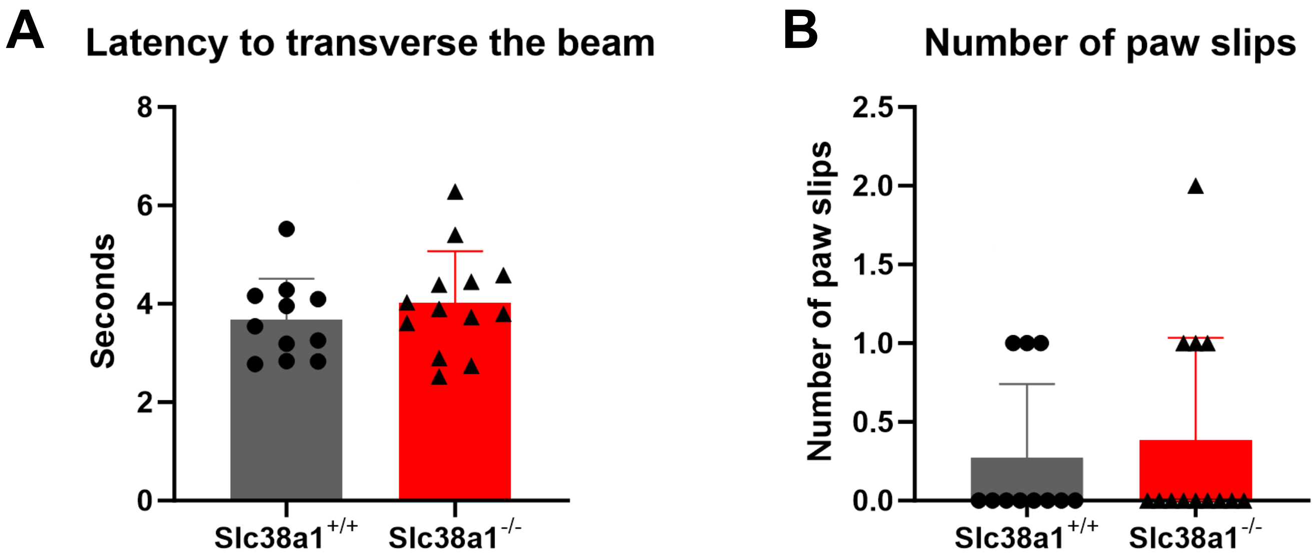


**Supplementary Figure 4. Slc38a1$^{-/-}$ mice do not have motor imbalance or miscoordination**
